# Pollen Ornamentation Morphospace: Interpretable Quantification by Persistent Homology

**DOI:** 10.64898/2026.11.18.767721

**Authors:** Qianzhi Deng, Keisuke Ishihara

## Abstract

Ornamentation, the sculpture of the outer pollen wall, is one of the central traits of pollen. However, it is mostly described with qualitative categories that rely on subjective judgment, and continuous, interpretable descriptors are still lacking. Here we used persistent homology, a fundamental method in topological data analysis, to define three interpretable descriptors (intensity, density and polarity) for three-dimensional pollen data. These descriptors form the three axes of a quantitative morphospace of pollen ornamentation, in which real, theoretical and ancestral pollen can be placed. The three axes were largely independent and separated common ornamentation types. In tests against known values, the morphospace recovered the number of ornamental elements in both theoretical pollen and real echinate pollen. Applied to pollination, it showed that, after phylogenetic correction, insect-pollinated pollen differed significantly from wind-pollinated pollen only in having more pronounced relief. In an evolutionary analysis within Asteraceae, it traced the three subfamilies diverging in different directions from their estimated common ancestor. By replacing qualitative categories with continuous, interpretable descriptors, this morphospace provides a common quantitative framework for pollen ornamentation. As three-dimensional pollen data accumulate, new grains can be placed in this morphospace without redefining the axes.

## 1 Introduction

Pollen is central to sexual reproduction in seed plants and is relevant to studies of pollinators, food security, human health and past climates (Hornick *et al*., 2022). Angiosperm pollen is highly diverse in morphology, varying markedly in size, shape, apertures and ornamentation (Halbritter *et al*., 2018). Size can be measured at high throughput by imaging flow cytometry (Hornick *et al*., 2025). Overall shape is conventionally summarized by the ratio of the polar axis to the equatorial diameter (P/E ratio; Punt *et al*., 2007) or quantified by elliptic Fourier descriptors of the outline (Bonhomme *et al*., 2013; Kriebel *et al*., 2017). Apertures are counted directly and represent one of the three characters of the NPC aperture classification system (Erdtman & Straka, 1961). In contrast, the characterization of ornamentation, the sculpture of the outer pollen wall, remains largely qualitative. Categories such as echinate (spiny), lophate (ridged), psilate (smooth) and reticulate (net-like) impose discrete boundaries on continuous variation and do not capture differences within each category. Assigning a grain to a category depends on subjective judgment, and the same grain may be classified differently (Halbritter *et al*., 2018). As a result, despite being one of the core traits of pollen, ornamentation still lacks a set of continuous, interpretable descriptors that would allow objective and reproducible comparisons.

Pollen ornamentation is difficult to quantify because it consists of many local elements, such as spines and ridges, whose form and number vary among grains. Existing morphometric methods are poorly suited to such structure. Fourier-based methods, whether elliptic Fourier analysis in two dimensions or spherical harmonic analysis in three dimensions, require the outline or surface to be closed (Kuhl & Giardina, 1982; Brechbühler *et al*., 1995), but the exine (pollen outer wall) can be interrupted by openings such as apertures. Even when the outline or surface is closed, a truncated Fourier or spherical harmonic series can produce spurious oscillations where the ornamentation changes abruptly (the Gibbs phenomenon; Gelb, 1997). Moreover, because each harmonic extends over the whole grain, the ornamentation is spread across many coefficients that are difficult to interpret biologically (Bookstein *et al*., 1982). Geometric morphometrics relies on landmarks that are homologous across specimens (Mitteroecker & Gunz, 2009), which are not available for pollen ornamentation.

Image-based methods are widely used to identify and compare pollen, based on features that are either handcrafted (Mander *et al*., 2013; Wei et al., 2025) or learned by machine learning (Romero *et al*., 2020). These features are often high-dimensional, and those learned by machine learning are usually hard to interpret (Barnes *et al*., 2023). Persistent homology, a fundamental method in topological data analysis, offers an alternative. Given a scalar field on a surface (e.g. the height of each point), persistent homology sweeps a threshold across the range of the field and records the thresholds at which each topological feature, such as a connected component, appears (its birth) and disappears (its death), with its lifetime known as its persistence (Edelsbrunner *et al*., 2002). LO-analysis, a classical light-microscopy method, examines ornamentation in a similar way by continuously varying the focus through the grain and recording the light patterns (Erdtman, 1956). Persistent homology requires no homologous landmarks, does not need the scalar field to be defined everywhere, and the persistence it records is stable under small perturbations of the field (Cohen-Steiner *et al*., 2007). While persistent homology has been proposed for quantifying pollen ornamentation (Li *et al*., 2018; Radja, 2021), it has yet to be applied to three-dimensional pollen data. Further, how to summarize the output of persistent homology as a result that is easily interpretable by a pollen biologist remains an open question.

Here we apply persistent homology to three-dimensional pollen data and introduce three interpretable descriptors: intensity *I*, density *D* and polarity *P*, which measure how pronounced the ornamentation is, how finely it is patterned and whether it is dominated by protrusions or by depressions, respectively. These three descriptors define a fixed morphospace that facilitates the comparison of theoretical, real and ancestral pollen without any free parameters. We first test the descriptors against known values: for theoretical pollen generated by a model of pattern formation (Radja *et al*., 2019), they recover the prescribed number of ornamental elements, and for real echinate pollen, they predict the number of spines. We then apply the morphospace to pollination, where, after phylogenetic correction, insectpollinated pollen differs from wind-pollinated pollen only in having more pronounced relief, and to evolution within Asteraceae, where the three subfamilies diverge in different directions from their estimated common ancestor. Together, we show that the three descriptors provide an interpretable quantification of pollen ornamentation, and that the resulting morphospace allows the study of pollen formation, function and evolution in a common quantitative framework.

## 2 Materials and Methods

### 2.1 Mapping pollen ornamentation to morphospace coordinates by persistent homology

Our pipeline maps each grain to its morphospace coordinates. Its input is a single-valued radial function *ρ* on the sphere, the distance from the center of the grain to its surface in each direction. In practice, *ρ* is sampled at the vertices of a grid formed by subdividing an icosahedron six times and is normalized by its maximum. Where *ρ* is undefined, the corresponding vertices are excluded together with their edges.

Sublevel-set filtrations of *ρ* and −*ρ* over this icosahedral grid give the depression and protrusion channels, respectively. We computed the zero-dimensional persistent homology of each channel (Edelsbrunner *et al*., 2002), discarded infinite or zero persistence, and pooled the remaining persistence of both channels into a set of signed persistence 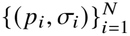, where *N* is the number of features, *pi* is the persistence of the *i*th feature, and *σi* is its channel, +1 for protrusion and −1 for depression. Persistence was computed with GUDHI 3.13.0 (Maria *et al*., 2014).

We define three scalars, intensity *I*, density *D* and polarity *P*, from the persistence fraction of each feature, *πi* = *pi*/ ∑_*j*_ *p j*, as

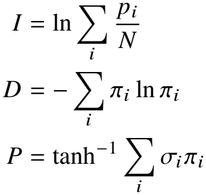

These three scalars form the axes of the morphospace, and their values for each grain are its morphospace coordinates.

### 2.2 Processing real pollen data from the 3D Pollen Library

We used three-dimensional meshes of real pollen from the 3D Pollen Library (Perry *et al*., 2017; Hayes, 2024), which were reconstructed from confocal images of exine autofluorescence and are openly available through NIH 3D. Ornamentation types of the specimens were taken from PalDat (Weber & Ulrich, 2017).

We restricted the analysis to near-spherical grains. Sphericity was defined as 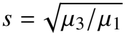, where *µ*1 ≥ *µ*2 ≥ *µ*3 are the eigenvalues of the covariance matrix of the mesh vertices. Of the 286 angiosperm pollen meshes, we retained the 162 whose sphericity exceeded 0.7. For each retained mesh, rays were cast from its center of mass along the directions given by the vertices of the icosahedral grid. The radial function *ρ* was taken as the distance to the farthest intersection, and directions without an intersection were omitted. The resulting *ρ* was then passed to the pipeline. Meshes were processed with trimesh 5.0.0 (Dawson-Haggerty et al., 2019) and rendered with PyVista 0.48.4 (Sullivan & Kaszynski, 2019).

Of the 162 grains retained for analysis, 20 were echinate Asteraceae. We excluded *Artemisia absinthium*, whose microechinate surface is dominated by three deep colpi rather than by spines, and manually counted the spines on the three-dimensional meshes of the remaining 19 grains to test the predicted number of spines.

### 2.3 Generating theoretical pollen from a model of pattern formation

Theoretical pollen grains are equilibrium states of the model of Radja *et al*. (2019), computed with the program released by the authors. In dimensionless form, the model is controlled by the wavenumber *l*0, the cubic coefficient *λ*3 and the reduced temperature *τ*. We varied *l*0 from 12.0 to 20.0 in steps of 0.5, set *λ*3 to −0.35, −0.25 and −0.15, and fixed *τ* at −0.25, giving 51 parameter sets. For each set, the program minimizes the free energy of the model and returns the equilibrium pattern as spherical harmonic coefficients. We reconstructed this pattern at the vertices of the icosahedral grid and passed the resulting values to the pipeline as *ρ*.

### 2.4 Testing the pollination syndrome and estimating ancestral ornamentation

Species names of the specimens were standardized to accepted names in WCVP (Govaerts *et al*., 2021) with rWCVP 1.3.0 (Brown *et al*., 2023). A dated phylogeny was built on the backbone of the GBOTB.extended.TPL megatree (Smith & Brown, 2018) with V.PhyloMaker2 0.1.0 (Jin & Qian, 2022). Species absent from the backbone were grafted onto their genus, and those whose genus was also absent were excluded. The morphospace coordinates of each species were averaged over its pollen grains, and all subsequent analyses were performed at the species level. Pollination modes were taken from the pollen vector records of the Ecological Flora of the British Isles (Fitter & Peat, 1994), in which a species can have more than one vector. Species names in this database were standardized as above and matched to the specimens by accepted name. Species recorded as insect but not wind were classed as insect-pollinated, and those recorded as wind but not insect as wind-pollinated; all other species were excluded. This left 116 species from 41 families, 95 insect-pollinated and 21 wind-pollinated.

Analyses of pollination mode used the dated phylogeny pruned to these 116 species. On each axis, the difference between insect- and wind-pollinated species was calculated as insect-pollinated minus wind-pollinated values. Without phylogenetic correction, the difference was expressed as Cohen’s *d*. With phylogenetic correction, it was estimated by phylogenetic generalized least squares (PGLS; Grafen, 1989), with Pagel’s *λ* (Pagel, 1999) fitted by maximum likelihood. Each PGLS difference was reported with its 95% confidence interval and tested with a *t*-test, and the *p*-values of the three axes were adjusted for multiple testing with the Benjamini–Hochberg procedure (Benjamini & Hochberg, 1995).

Analyses within Asteraceae used the dated phylogeny pruned to the 28 Asteraceae species, regardless of pollination records. On each axis, phylogenetic signal was measured by Pagel’s *λ*, estimated by maximum likelihood with a 95% confidence interval from its likelihood profile. Each *λ* was tested against *λ* = 0 with a likelihood-ratio test, and the *p*-values of the three axes were adjusted in the same way as above. Ancestral values on each axis were estimated by maximum likelihood under a Brownian motion model (Schluter *et al*., 1997).

All phylogenetic analyses were run in R 4.4.3 (R Core Team, 2025). Trees were handled with ape 5.8.1 (Paradis & Schliep, 2019). PGLS of pollination mode was fitted with the gls function of nlme 3.1-170 (Pinheiro *et al*., 2026). Within Asteraceae, Pagel’s *λ* was fitted with caper 1.0.4 (Orme *et al*., 2025), and ancestral states were estimated with the fastAnc function of phytools 2.5.2 (Revell, 2024).

## 3 Results

### 3.1 Persistent homology decomposes pollen ornamentation into three interpretable axes

We illustrate how persistent homology captures ornamentation using a schematic grain surface *ρ* bearing three protrusions and three depressions (Fig. 1a). Consider a spherical water surface of radius *r* centered on the grain’s center of mass, and let *r* sweep across the range of *ρ*. The sweep defines a filtration, with *r* as the filtration value. The part of the pollen surface under water, {*ρ* ≤ *r*}, is the sublevel set of *ρ* at *r*, and forms the depression channel; the part above water, {*ρ* ≥ *r*}, is equivalently the sublevel set of -*ρ* at -*r*, and forms the protrusion channel. In the depression channel, rising water forms a pool in each depression; in the protrusion channel, falling water exposes an island at each protrusion. Each pool or island disappears when it merges with an older one at a saddle. At each filtration value, the reduced zero-dimensional homology counts the separate pools 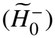 or islands 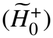 except the oldest pool or island, which never disappears. For example, three separate pools at *r* = 0.5 give 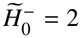, and three separate islands at *r* = 0.75 give 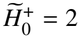. Persistent homology then follows each finite component from its birth to death, giving a persistence bar with persistence *pi* (the depth of a depression or the height of a protrusion) and channel *σi* (+1 for protrusion and −1 for depression). Together the bars form a barcode of signed persistence, and the number of finite bars crossing any filtration value equals 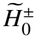 at that value (Fig. 1b, top). Plotting the death of each component against its birth gives the persistence diagram (Fig. 1b, bottom).

**Figure 1.**
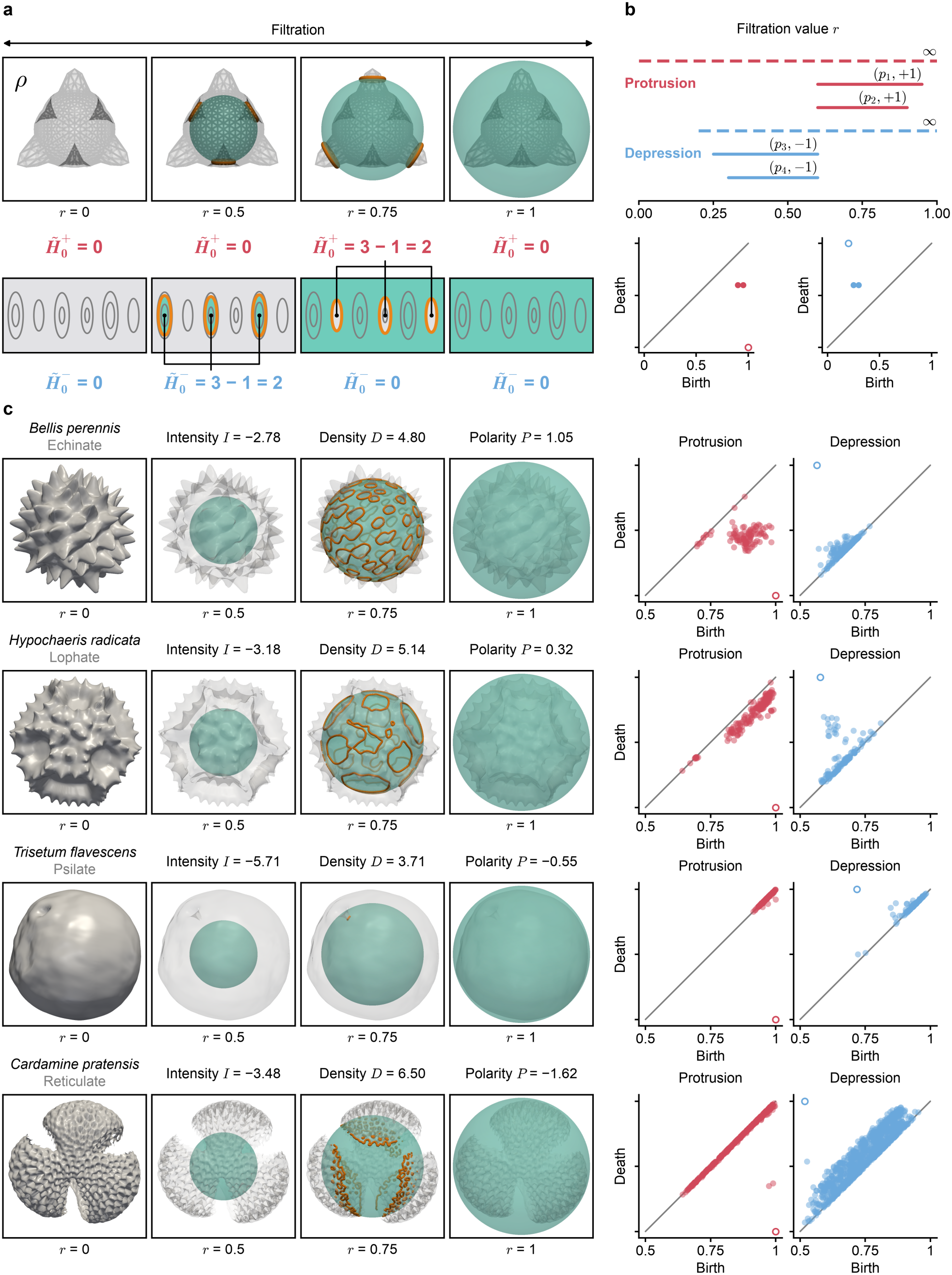
Signed persistence of pollen ornamentation. (a) A schematic grain *ρ* with three protrusions and three depressions at filtration values *r* = 0, 0.5, 0.75 and 1, shown in three dimensions (top) and as an unrolled equatorial band (bottom), with 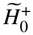 and 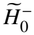 at each value. Teal, water surface; orange, shoreline (*ρ* = *r*). (b) Persistence barcode (top) and persistence diagrams (bottom) of the grain in (a). Red, protrusion channel; blue, depression channel; dashed bars and open circles, essential components excluded from the analysis. (c) Four real grains, one per ornamentation type: *Bellis perennis* (3DPX-017818, echinate), *Hypochaeris radicata* (3DPX-021466, lophate), *Trisetum flavescens* (3DPX-020938, psilate) and *Cardamine pratensis* (3DPX-020611, reticulate). Left, the exine and water surface at the filtration values in (a), with the intensity *I*, density *D* and polarity *P* of each grain; right, persistence diagrams. Colors and symbols as in (a) and (b).

The three descriptors defined from signed persistence (see Materials and Methods) can be interpreted both in the persistence diagram and on the pollen surface (Fig. 1c). Intensity *I*, the logarithm of the mean persistence, measures how far the points lie from the diagonal on average and hence the mean amplitude of the ornamental relief. Among the four grains, *I* is highest in the echinate pollen of *Bellis perennis* (*I* = −2.78), whose tall spines give high persistence, slightly lower in the lophate pollen of *Hypochaeris radicata* (*I* = −3.18), whose spines are shorter than those of *B. perennis*, and lowest in the psilate pollen of *Trisetum flavescens* (*I* = −5.71), whose shallow relief gives low persistence. Density *D*, the Shannon entropy of the persistence fractions, reflects how evenly the persistence is spread, that is, how finely the ornamentation is patterned.

*D* is highest in the reticulate pollen of *Cardamine pratensis* (*D* = 6.50), whose numerous lumina form a large cloud of points in the depression channel, and lowest in *T. flavescens* (*D* = 3.71), whose smooth surface is barely patterned. Polarity *P* compares the fractions of persistence carried by the two channels and thereby indicates whether the ornamentation is dominated by protrusions or by depressions. *P* is high in *B. perennis* (*P* = 1.05), whose spines carry most of the persistence, lower in *H. radicata* (*P* = 0.32), where the lacunae between ridges add persistence to the depression channel, and strongly negative in *C. pratensis* (*P* = −1.62), whose lumina carry most of the persistence in the depression channel. Each descriptor therefore corresponds to a feature of ornamentation that can be recognized by eye.

We placed all 162 near-spherical angiosperm grains in the morphospace (Fig. 2). They spread across all three axes, with intensity *I* ranging from −7.03 to −2.68, density *D* from 1.11 to 7.46 and polarity *P* from −1.74 to 1.41. The axes were only weakly correlated, with Pearson correlation coefficients of −0.15 between *I* and *D*, 0.18 between *I* and *P* and 0.09 between *D* and *P*. Five example grains of each common ornamentation type illustrate where these types lie in the morphospace: the echinate grains at high *I* and high *P*, the lophate grains at high *I* and lower *P*, the psilate grains at low *I* and low *D* and the reticulate grains at negative *P* and high *D*. Overall, persistent homology decomposes pollen ornamentation into three largely independent, interpretable axes, and the common ornamentation types fall in different regions of the space they span.

**Figure 2.**
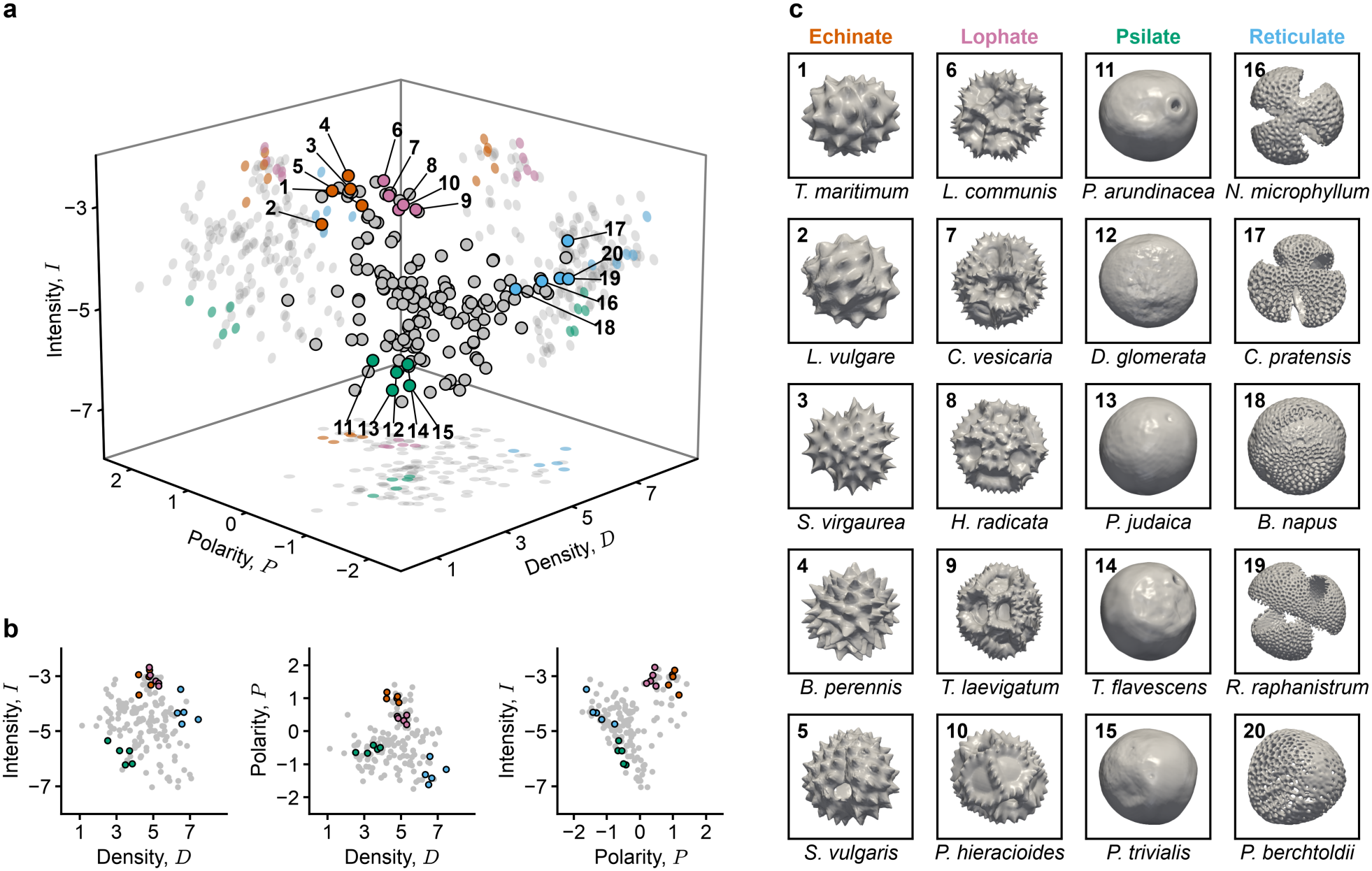
Real pollen in the morphospace. (a) The 162 near-spherical angiosperm grains in the morphospace of intensity *I*, density *D* and polarity *P*. Gray, all grains; colored and numbered, the representative grains in (c). (b) The same grains on the *D*–*I, D*–*P* and *P*–*I* faces. Colors as in (a). (c) Twenty representative grains, five per common ornamentation type. Echinate: 1, *Tripleurospermum maritimum* (3DPX-021473); 2, *Leucanthemum vulgare* (3DPX-017803); 3, *Solidago virgaurea* (3DPX-021264); 4, *Bellis perennis* (3DPX-017818); 5, *Senecio vulgaris* (3DPX-020862). Lophate: 6, *Lapsana communis* (3DPX-022934); 7, *Crepis vesicaria* subsp. *taraxacifolia* (3DPX-023360); 8, *Hypochaeris radicata* (3DPX-021466); 9, *Taraxacum laevigatum* (3DPX-021472); 10, *Picris hieracioides* (3DPX-021105). Psilate: 11, *Phalaris arundinacea* (3DPX-021112); 12, *Dactylis glomerata* (3DPX-022933); 13, *Parietaria judaica* (3DPX-021111); 14, *Trisetum flavescens* (3DPX-020938); 15, *Poa trivialis* (3DPX-022883). Reticulate: 16, *Nasturtium microphyllum* (3DPX-021959); 17, *Cardamine pratensis* (3DPX-020611); 18, *Brassica napus* (3DPX-017819); 19, *Raphanus raphanistrum* (3DPX-023493); 20, *Potamogeton berchtoldii* (3DPX-022903).

### 3.2 Morphospace coordinates recover the number of ornamental elements

To quantitatively assess the validity of the morphospace, we analyzed pollen whose number of ornamental elements is known. Radja et al. (2019) developed a physical model where pollen ornamentation forms as the primexine (a polysaccharide layer that templates the exine) phase-separates into regularly spaced elements. In this model, the wavenumber *l*0 specifies how many times the elements repeat around the grain and thus sets the number of ornamental elements. Using this model, we generated theoretical pollen of varying *l*0 and projected the resulting patterns onto the morphospace (Fig. 3a). We asked whether *l*0 can be recovered from *D* and *P*.

**Figure 3.**
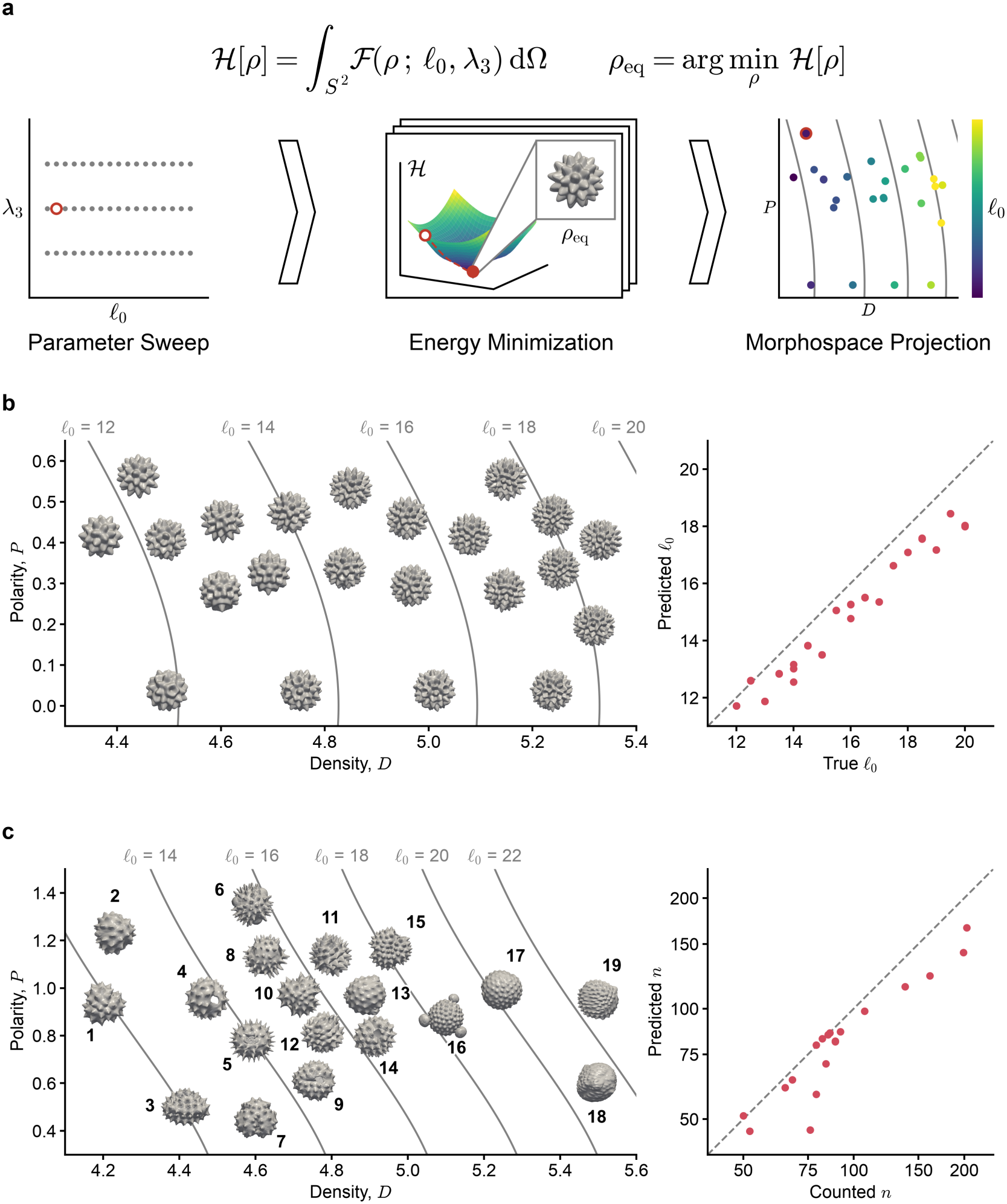
Recovery of the number of ornamental elements. (a) Generation of theoretical pollen with the model of Radja et al. (2019): parameter sweep over *l*0 and *λ*3 (left), energy minimization to the equilibrium *ρ*eq (center) and projection onto the morphospace *D*–*P* face (right). Red, one parameter set followed through the three steps; gray lines, iso-*l*0 lines; the energy surface is schematic. (b) Theoretical pollen on the *D*–*P* face (left; coincident grains drawn once) and *l*0 predicted from *D* and *P* against the true value (right). The relief of theoretical pollen is schematic, as the model does not specify the height of ornamentation. Gray lines as in (a). (c) The 19 echinate Asteraceae pollen grains in Supporting Information B on the *D*–*P* face (left) and spine number *n* predicted from *D* and *P* against the counted number (right; logarithmic axes). Lines as in (b).

We derived that *D* and *P* are related to the wavenumber *l*_0_ as

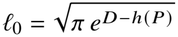

where *h* (*P*) is the binary entropy of the persistence fractions of the two channels (Supporting Information A). Since the elements tile the whole surface, a pattern with *l*0 elements around the grain has about *l*^2^_0_ *π* elements in each channel, so the same relation also predicts the number of spines *n* on real echinate pollen as

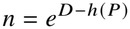

For theoretical pollen, the predicted *l*0 differed from the true value by 6.1% on average (Fig. 3b). We further tested our prediction for real pollen, where we manually counted the spines on the three-dimensional meshes of 19 echinate Asteraceae grains (see Materials and Methods). These grains bore 50 to 203 spines each, and the predicted number differed from the count by 11.9% on average (Fig. 3c; Supporting Information B). *D* and *P* thus recover the number of ornamental elements in both theoretical and real pollen.

### 3.3 The intensity axis reflects the classical pollination syndrome

A quantitative morphospace makes it possible to test qualitative theories in biology (Chartier *et al*., 2014). We test the pollination syndrome, which posits that plants pollinated by the same vector, such as wind or insects, converge on similar floral traits. Flowers of wind-pollinated plants, for example, tend to be small and inconspicuous and lack nectaries (Friedman & Barrett, 2009). For pollen morphology, the syndrome predicts that wind-pollinated pollen is smoother, which lowers its settling velocity and lets it travel farther on the wind, whereas insect-pollinated pollen bears more conspicuous ornamentation, which helps it attach to pollinators (Rosbakh *et al*., 2026). Testing these predictions across the Central European flora, the same study scored ornamentation types as present or absent and found that only reticulate ornamentation differed between wind- and insect-pollinated pollen. Using our quantitative morphospace, we instead measure how much windand insect-pollinated pollen differ along the three axes of intensity, density and polarity.

We compared 95 insect-pollinated and 21 wind-pollinated species on their dated phylogeny (Fig. 4a). Without phylogenetic correction, insect-pollinated pollen had higher values than wind-pollinated pollen on all three axes (Fig. 4b), with Cohen’s *d* of 1.10 for *I*, 0.72 for *D* and 0.42 for *P*. After phylogenetic correction by PGLS (Fig. 4c), the difference remained significant only for *I* (0.87, 95% CI 0.44 to 1.30, *p*FDR < 0.001), not for *D* (0.49, 95% CI −0.15 to 1.13, *p*FDR = 0.129) or *P* (0.23, 95% CI −0.04 to 0.51, *p*FDR = 0.129). Wind-pollinated pollen therefore has lower relief than insect-pollinated pollen, as the pollination syndrome predicts, but does not differ significantly in the fineness of the ornamentation or in whether it is dominated by protrusions or by depressions.

**Figure 4.**
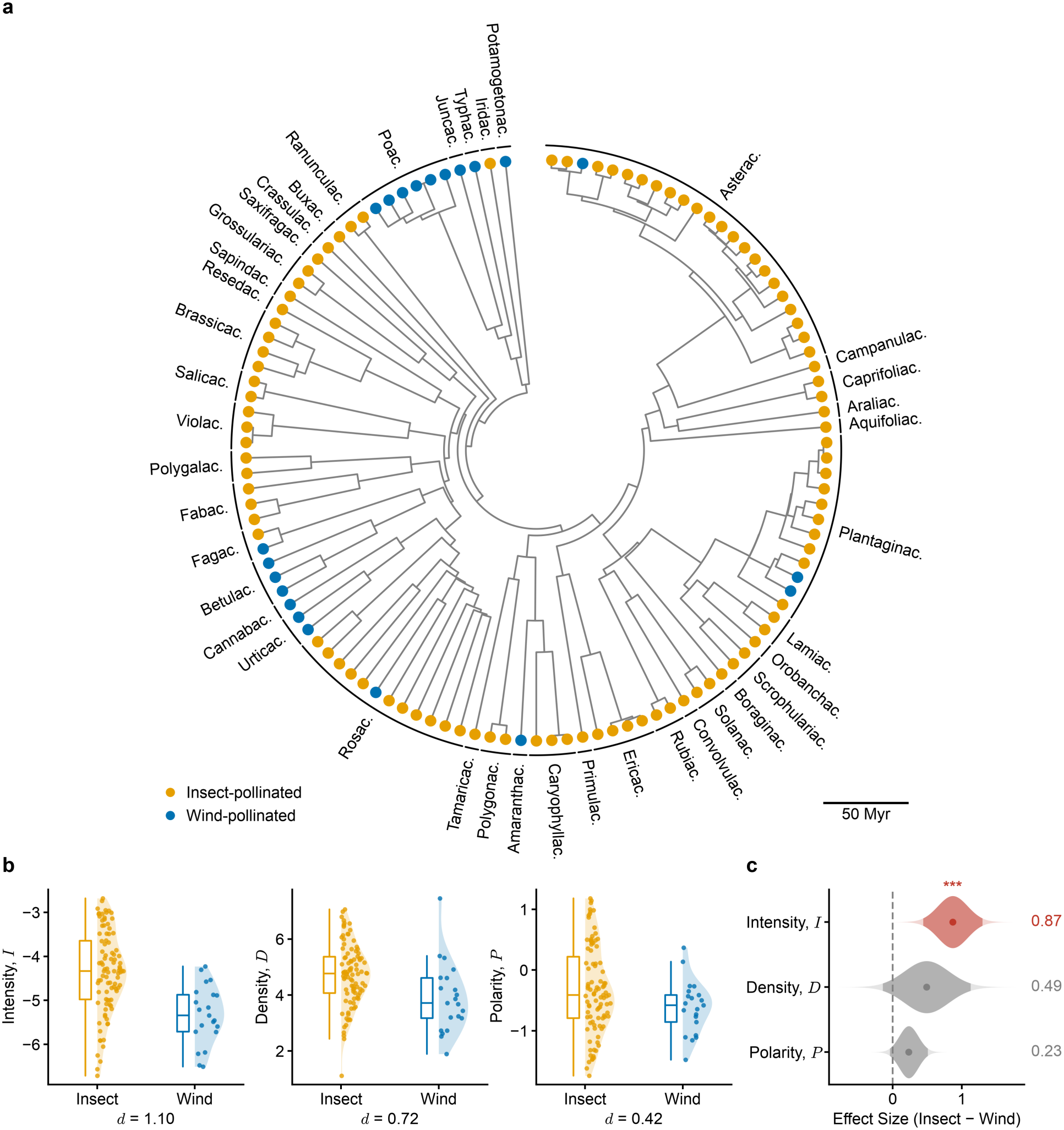
Insect- and wind-pollinated pollen on the three axes of the morphospace. (a) Dated phylogeny of the 116 species from 41 families, with pollination modes at the tips and abbreviated family names on the outer arcs. Yellow, insect-pollinated; blue, wind-pollinated. Scale bar, 50 Myr. (b) Intensity, density and polarity of insect- and wind-pollinated species with Cohen’s *d* reported; colors as in (a). (c) Differences between the groups (insect − wind) estimated by PGLS, with values on the right. Violins, *t* distributions of the estimates with 95% confidence intervals; asterisks, *p*FDR < 0.05 (*), < 0.01 (**) and < 0.001 (***).

### 3.4 Asteraceae subfamilies show evolutionary divergence in the morphospace

The evolution of pollen morphology is increasingly studied in phylomorphospaces, in which species and their ancestors are placed together in a morphospace (Jardine *et al*., 2022; Luo *et al*., 2026). In these studies, ornamentation is scored as discrete characters and the morphospace is obtained by ordination of these characters. A change in ornamentation therefore appears as a switch between character states. Our morphospace measures ornamentation along continuous axes, so it can also show in which directions, and how far, ornamentation has evolved. As an example, we traced these changes in Asteraceae, the family best represented in our dataset.

We placed the 28 Asteraceae species in the morphospace together with their ancestral states, estimated on each axis under Brownian motion (Fig. 5a,b). From the estimated root, the three subfamilies shifted in different directions. Cichorioideae shifted toward higher density and lower polarity, that is, toward finer and more porous ornamentation, as expected from their lophate pollen (Jardine *et al*., 2022). Asteroideae shifted the opposite way, toward coarser or-namentation dominated by protrusions, and Carduoideae shifted toward lower intensity, that is, toward lower relief. *Artemisia absinthium*, the only wind-pollinated species among the 23 with pollination records, lay apart from the rest at the low end of all three axes. This is consistent with the smaller spines of *Artemisia* pollen (Jardine *et al*., 2022) and with the characters of wind-pollinated pollen in the previous section. In this way, the morphospace turns the evolution of ornamentation into movement along interpretable axes.

**Figure 5.**
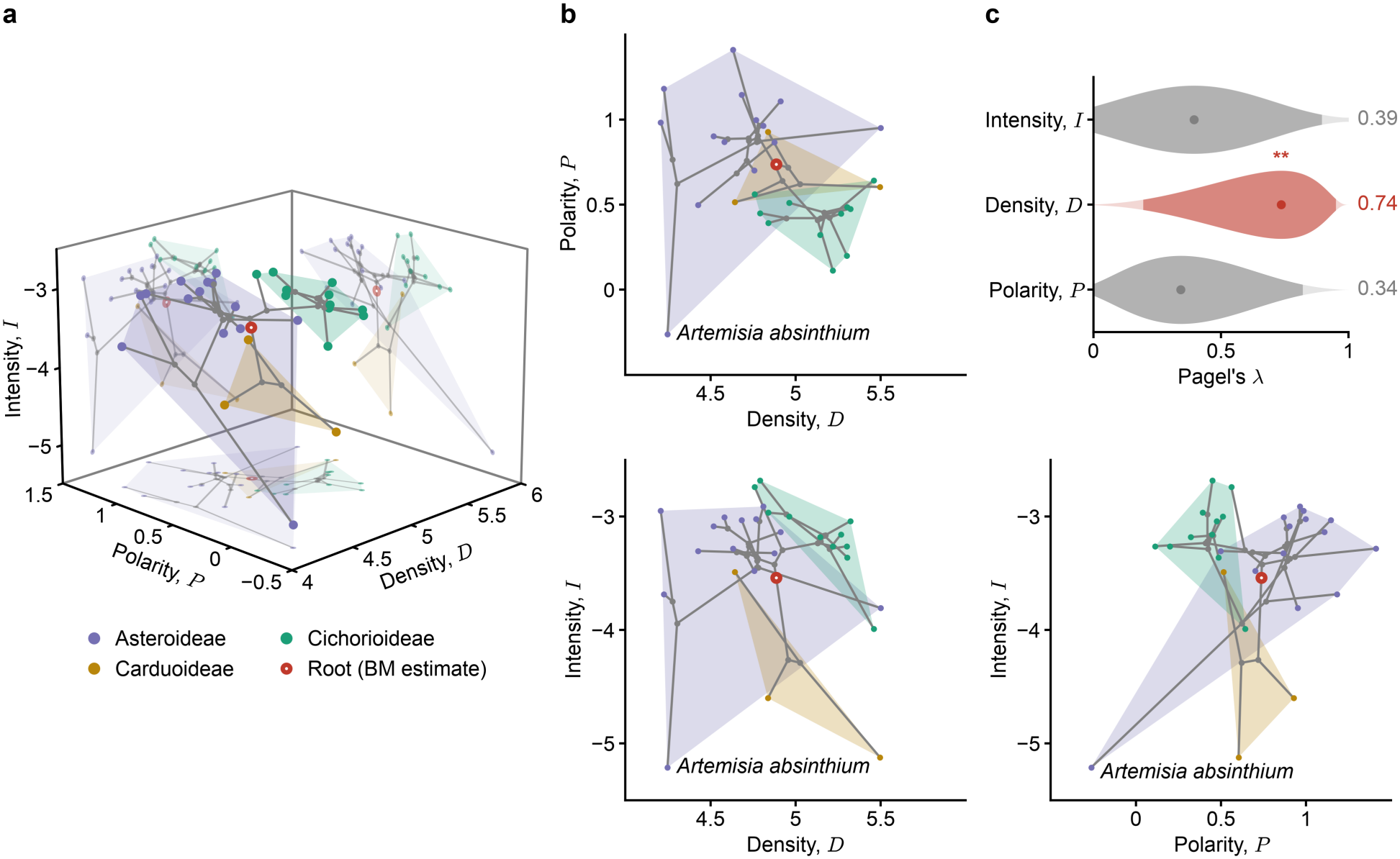
Evolution of pollen ornamentation in Asteraceae in the morphospace. (a) Phylomorphospace of the 28 species. Extant species are colored by subfamily, and the shaded areas enclose the species of each subfamily. Gray lines and dots show the branches and estimated ancestral states, and the red ring marks the estimated root. (b) The phylomorphospace on its *D*–*P, D*–*I* and *P*–*I* faces, with *Artemisia absinthium* labeled. Colors and symbols as in (a). (c) Pagel’s *λ* for the three axes, with values on the right. Violins, profile likelihoods with 95% confidence intervals; asterisks as in Fig. 4.

To assess how closely each axis follows the phylogeny, we estimated Pagel’s *λ* within Asteraceae (Fig. 5c). Only *D* showed a significant phylogenetic signal (*λ* = 0.74, 95% CI 0.19 to 0.95, *p*FDR = 0.007). The signal in *I* (*λ* = 0.39, 95% CI 0 to 0.90, *p*FDR = 0.164) and *P* (*λ* = 0.34, 95% CI 0 to 0.83, *p*FDR = 0.090) was not significant. Of the three axes, density thus tracks the phylogeny most closely, as it separates the echinate pollen of Asteroideae from the lophate pollen of many Cichorioideae.

## 4 Discussion

We have used persistent homology to construct a quantitative morphospace for pollen ornamentation, whose interpretable axes allow theoretical, real and ancestral pollen to be placed together. Morphospaces are either the-oretical, with axes defined in advance, or empirical, with axes obtained from an ordination of specimens (McGhee, 1999). The classic example of a theoretical morphospace is that of shell coiling by Raup (1966), which founded theoretical morphology and originated in his attempt to capture the “spectra of gradation” between qualitative categories such as turriculate and turbiniform (Raup, 1961). Our morphospace pursues a similar aim for pollen ornamentation, and its axes are likewise defined in advance. Beyond this, the axes have no free parameters and are computed fully automatically, so the coordinates require neither arbitrary parameter choices nor manual intervention. The axes can also be interpreted directly as the intensity, density and polarity of the ornamentation, whereas the axes of empirical morphospaces have only statistical meaning and must be interpreted post hoc (Mitteroecker & Gunz, 2009). Finally, since the three axes capture distinct aspects of ornamentation, differences along each can be analyzed separately. By contrast, the independence of the axes of classical morphospaces has long been debated; even in Raup’s space, the axes covary mathematically (Schindel, 1990).

Our approach can be viewed as a generalization of LO-analysis. LO-analysis follows changes in the brightness of the ornamentation as the focus moves up and down. As the focal plane descends, protrusions appear first bright and then dark (the LO-pattern), while depressions appear first dark and then bright (the OL-pattern; Erdtman, 1956). In our approach, the filtration value *r* corresponds geo-metrically to a spherical water surface of radius *r*, which for near-spherical pollen locally approximates the focal plane. When the water level falls, the protrusions emerge one by one as islands, and the water breaks into pools that retreat into the depressions. These two processes mirror the LO- and OL-patterns. As a generalization, persistent homology extends LO-analysis in two respects. It covers all filtration values rather than a few focal planes, and its results are quantitative and do not depend on the judgment of an observer.

One limitation of this study is the size and taxonomic coverage of our sample. Because palynological records consist mainly of two-dimensional light and scanning electron micrographs, few three-dimensional pollen data exist beyond the 3D Pollen Library used here, which is the largest public collection to date. Further limitations arise from the pipeline itself. First, the coordinates may be sensitive to differences in mesh resolution. Resampling all meshes to the same resolution might reduce this effect. Second, the pipeline assumes the input pollen to be near-spherical, so for markedly non-spherical pollen, the overall shape could be mistaken for ornamentation. For example, the poles of an ellipsoidal pollen grain would be read as long spines. A possible alternative would be to measure ornamentation relative to the overall shape, such as that given by a low-order spherical harmonic expansion, rather than to a sphere. Third, the pipeline treats apertures as ornamental depressions, so both enter the depression channel. The effect of this conflation is relatively limited, however, because the three axes describe statistical features of all persistence bars, and apertures, in most cases, account for only a small proportion of them. Fourth, to keep the framework simple, we based our morphospace only on zero-dimensional persistent homology, which does not record the shape or spatial arrangement of ornamental elements. The morphospace therefore cannot distinguish ornamentation that differs only in these respects. Introducing higher-dimensional homology may overcome this limitation.

Pollen ornamentation has long been described by qualitative categories. Our three scalars based on persistent homology offer a continuous, interpretable quantification, and new grains can be placed directly in the morphospace they span. As larger and more balanced samples of three-dimensional pollen data accumulate, this morphospace may provide a common framework for pollen ornamentation.

## Acknowledgements

We thank Anthony J. Hayes and the Bioimaging Research Hub, Cardiff University, for generously making the 3D Pollen Library available to the scientific community, with-out which this study would not have been possible. We thank members of the Ishihara lab, in particular, Yajushi Khurana, for comments on the manuscript. Claude Opus 4.8 (Anthropic) was used to assist with code development. All AI-assisted outputs were reviewed, edited and validated by the authors. This research received no specific funding.

## Author contributions

QD was involved in conceptualization, methodology, software, formal analysis, investigation, data curation, visualization and writing – original draft, review and editing. KI was involved in supervision and writing – review and editing.

## Data availability

The three-dimensional pollen data analyzed in this study are openly available from the 3D Pollen Library on NIH 3D (https://3d.nih.gov/collections/33). Pollination modes were obtained from the Ecological Flora of the British Isles (http://ecoflora.org.uk), and ornamentation types from PalDat (https://www.paldat.org). The code and data needed to reproduce all analyses are available on GitHub (https://github.com/qianzhi-deng/Pollen-Ornamentation-Morphospace).

## Supporting Information

### A Derivation of the relation between density, polarity and wavenumber

By definition, *D* is the Shannon entropy of the persistence fractions *πi*. If the ornamentation has only one channel and consists of *n* elements of equal persistence, each element has the fraction *πi* = 1/*n*, so *D* = ln *n*, exactly the logarithm of the number of elements. Ornamentation, however, usually has both protrusions and depressions. To account for both channels, we introduce the binary entropy of their fractions.

Write the fractions of the protrusion and depression channels as

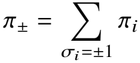

The two fractions sum to 1, and by the definition of *P* their difference is *π*+ − *π*− = _*i*_ *πiσi* = tanh *P*. Hence

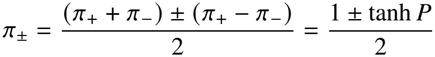

and their binary entropy is

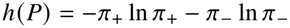

If each channel consists of *n* elements of equal persistence, each element in the protrusion or depression channel has the fraction *π*+/*n* or *π*−/*n*. Substituting into the definition of *D* gives

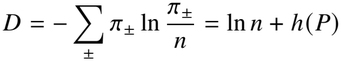

Since *l*0 counts how many times the pattern repeats around the grain, *l*0 = 2*πR*/*λ*, where *R* is the radius of the grain and *λ* is the characteristic wavelength of the pattern. If each period of the pattern occupies an area of *λ*^2^, each channel contains 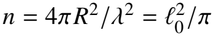 elements, so

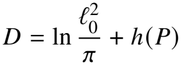

Rearranging gives

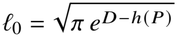

### B Counted and predicted spine numbers of 19 echinate Asteraceae pollen grains

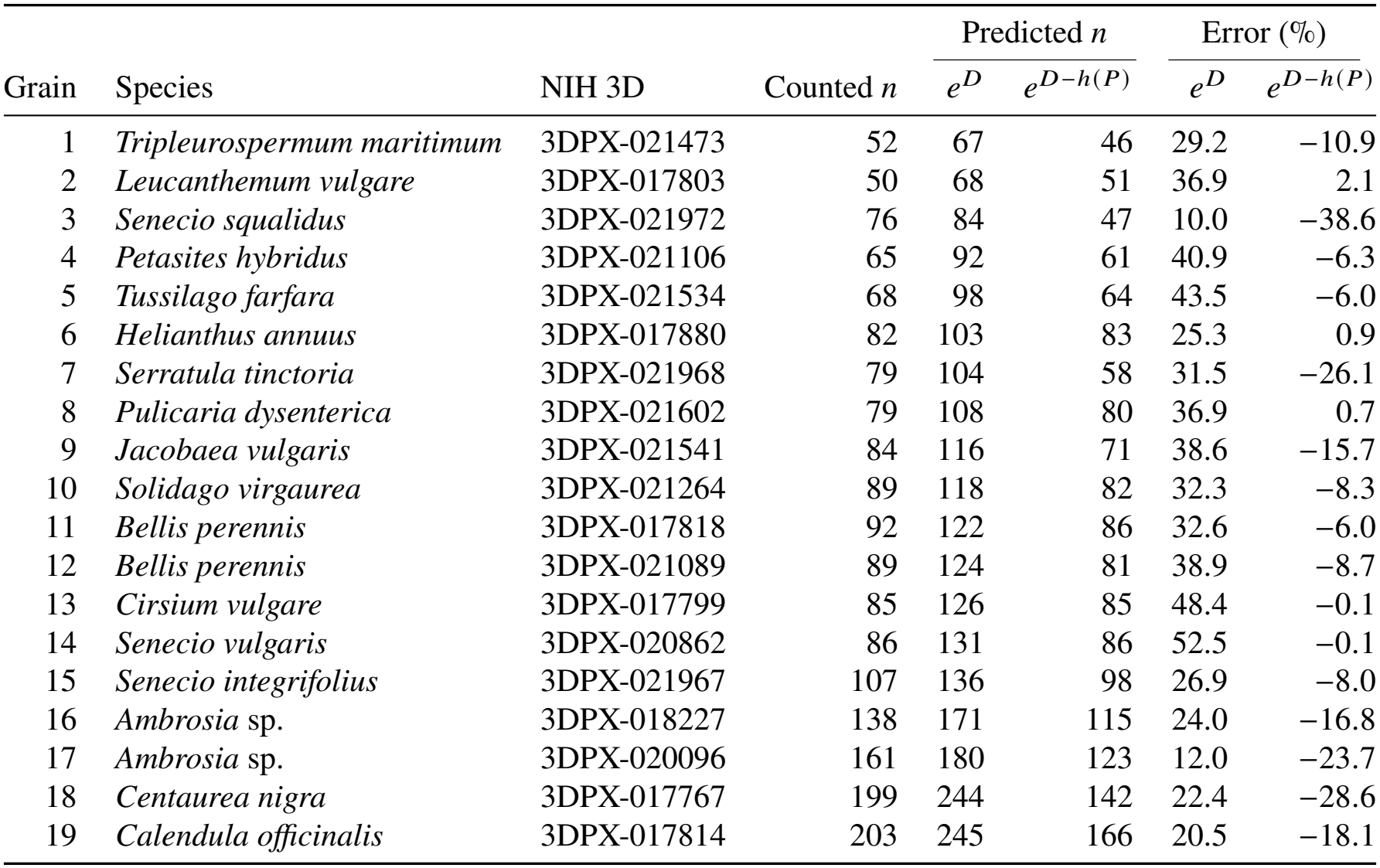

| Grain | Species | NIH 3D | Counted *n* | Predicted *n* |  | Error (%) |  |
|---|---|---|---|---|---|---|---|
|  |  |  |  | *e*<sup>*D*</sup> | *e*<sup>*D*−*h*(*P*)</sup> | *e*<sup>*D*</sup> | *e*<sup>*D*−*h*(*P*)</sup> |
| 1 | *Tripleurospermum maritimum* | 3DPX-021473 | 52 | 67 | 46 | 29.2 | −10.9 |
| 2 | *Leucanthemum vulgare* | 3DPX-017803 | 50 | 68 | 51 | 36.9 | 2.1 |
| 3 | *Senecio squalidus* | 3DPX-021972 | 76 | 84 | 47 | 10.0 | −38.6 |
| 4 | *Petasites hybridus* | 3DPX-021106 | 65 | 92 | 61 | 40.9 | −6.3 |
| 5 | *Tussilago farfara* | 3DPX-021534 | 68 | 98 | 64 | 43.5 | −6.0 |
| 6 | *Helianthus annuus* | 3DPX-017880 | 82 | 103 | 83 | 25.3 | 0.9 |
| 7 | *Serratula tinctoria* | 3DPX-021968 | 79 | 104 | 58 | 31.5 | −26.1 |
| 8 | *Pulicaria dysenterica* | 3DPX-021602 | 79 | 108 | 80 | 36.9 | 0.7 |
| 9 | *Jacobaea vulgaris* | 3DPX-021541 | 84 | 116 | 71 | 38.6 | −15.7 |
| 10 | *Solidago virgaurea* | 3DPX-021264 | 89 | 118 | 82 | 32.3 | −8.3 |
| 11 | *Bellis perennis* | 3DPX-017818 | 92 | 122 | 86 | 32.6 | −6.0 |
| 12 | *Bellis perennis* | 3DPX-021089 | 89 | 124 | 81 | 38.9 | −8.7 |
| 13 | *Cirsium vulgare* | 3DPX-017799 | 85 | 126 | 85 | 48.4 | −0.1 |
| 14 | *Senecio vulgaris* | 3DPX-020862 | 86 | 131 | 86 | 52.5 | −0.1 |
| 15 | *Senecio integrifolius* | 3DPX-021967 | 107 | 136 | 98 | 26.9 | −8.0 |
| 16 | *Ambrosia* sp. | 3DPX-018227 | 138 | 171 | 115 | 24.0 | −16.8 |
| 17 | *Ambrosia* sp. | 3DPX-020096 | 161 | 180 | 123 | 12.0 | −23.7 |
| 18 | *Centaurea nigra* | 3DPX-017767 | 199 | 244 | 142 | 22.4 | −28.6 |
| 19 | *Calendula officinalis* | 3DPX-017814 | 203 | 245 | 166 | 20.5 | −18.1 |

